# The Energetic Cost of Building Human Skeletal Muscle

**DOI:** 10.64898/2026.08.17.745156

**Authors:** Michalis G. Nikolaidis, Vassilis Paschalis, Nikos V. Margaritelis

## Abstract

The energetic cost of building human skeletal muscle has never been explicitly calculated or measured. We developed a quantitative bottom-up accounting model that integrates human skeletal-muscle composition with empirically informed estimates of tissue synthesis, physiological deposition, maintenance during accretion, and diet-induced thermogenesis. The calculation was expressed per kg of wet skeletal muscle and organized into five additive components: stored tissue energy, biochemical synthesis cost, physiological deposition cost, resting maintenance during accretion, and diet-induced thermogenesis. Stored tissue energy was approximately 5670 kJ/kg (1355 kcal/kg). Adding biochemical synthesis cost gave 6340 kJ/kg (1515 kcal/kg). Applying empirically derived deposition-efficiency parameters yielded a physiological deposition requirement of 9780 to 11690 kJ/kg (2338 to 2793 kcal/kg), centrally 10830 kJ/kg (2587 kcal/kg). Adding resting maintenance during accretion and diet-induced thermogenesis produced a final additional metabolizable energy intake of 13410 to 15520 kJ/kg (3204 to 3710 kcal/kg), centrally 14570 kJ/kg (3481 kcal/kg). This value provides a first quantitative reference estimate for the energetic cost of human skeletal-muscle accretion.

## 1. INTRODUCTION

Skeletal muscle is central to human physiology, nutrition, rehabilitation, aging, and medicine. Athletes seek to increase it, older adults try to preserve it, and patients recovering from disease or functional decline may need to regain it. Yet a basic quantitative question remains unresolved: what is the energetic cost of building human skeletal muscle? The energetic requirements of hypertrophy, realistic rates of fat-free mass accretion, and the rationale for dietary energy surpluses have been discussed previously (Slater et al. 2019; Larson-Meyer et al. 2022; Trexler et al. 2026), but the energetic cost of constructing skeletal-muscle tissue itself has not been directly measured in adult humans, and no integrated tissue-specific estimate currently accounts explicitly for its major energetic components.

This gap has both methodological and practical consequences. In research design and applied practice, expected muscle gain, energy prescription, intervention duration, and outcome interpretation should be physiologically compatible. If a researcher, clinician, dietitian, or coach expects approximately 1 kg of skeletal-muscle gain, it should be possible to judge whether the associated energetic assumptions are biologically plausible. Without such a reference point, several related but distinct quantities can easily be conflated: the chemical energy stored in newly formed tissue, the biochemical cost of synthesizing its constituents, the broader physiological cost of tissue deposition, the energy required to maintain newly accumulated tissue during the period of accretion, the additional dietary energy that may be required to supply these processes, and practical recommendations for daily energy surplus. These represent distinct levels of energetic accounting and should therefore be distinguished explicitly.

We therefore approached the problem using first-principles reasoning supported by empirical constraints. In effect, the analysis adopts a Fermi-style strategy to a quantity that has likely remained poorly defined not because it is unimportant, but because it is biologically composite and difficult to measure directly in humans. Rather than treating this complexity as a reason to leave the quantity unspecified, we decompose the problem into physiologically interpretable components and use available empirical evidence to estimate its expected order of magnitude. This bottom-up approach is consistent with quantitative treatments of tissue deposition in which stored energy and the energetic costs of deposition are considered separately (Hall 2010), while allowing the assumptions contributing to the final estimate to remain explicit and testable.

The present analysis therefore develops a quantitative bottom-up accounting model for the energetic cost of building 1 kg of wet human skeletal muscle. The calculation is expressed per kg of wet mixed skeletal muscle and integrates empirical estimates of muscle composition with energetic parameters describing tissue synthesis, physiological deposition, maintenance during accretion, and diet-induced thermogenesis. The model is organized into five components: stored tissue energy, biochemical synthesis cost, physiological deposition cost, resting maintenance during accretion, and diet-induced thermogenesis. The purpose is not to define an invariant biological constant or a prescriptive daily energy surplus, but to provide a transparent, tissue-specific quantitative reference estimate against which claims concerning skeletal-muscle accretion and its energetic requirements can be evaluated.

## 2. METHODS AND ASSUMPTIONS

### 2.1 Reference tissue and composition

The calculation is expressed per 1 kg of wet human mixed skeletal muscle. This unit is not intended to represent a universal biological target, but to provide a practical denominator for estimating and comparing the energetic costs of skeletal-muscle accretion.

The reference composition was: protein, 177 g/kg wet muscle (Stroh et al. 2021); lipid, 30 g/kg wet muscle (Goodpaster et al. 2004); and glycogen, 20 g/kg wet muscle (Jensen et al. 2011). DNA, RNA, free amino acids, lactate, minerals, inorganic ions, and other minor constituents were not included as separate energy-retaining pools because they are quantitatively small or contribute little or no retained chemical energy in this accounting. Creatine and phosphocreatine were also not included as separate energy-retaining pools. Although 1 kg of wet skeletal muscle contains approximately 1.3 g free creatine and 4.3 g phosphocreatine (Bonilla et al. 2021), the energetic cost of synthesizing this creatine pool and phosphorylating the phosphocreatine fraction is approximately 6.5 kJ/kg wet muscle (1.6 kcal/kg). This is negligible relative to the retained energy in protein, lipid, and glycogen and does not affect the rounded estimate.

### 2.2 Stored tissue energy parameters

The gross-energy values used to calculate stored tissue energy were 23.5 kJ/g (5.62 kcal/g) for protein (Southgate and Durnin 1970; Reeds et al. 1980), 38.9 kJ/g (9.30 kcal/g) for lipid (Whyte et al. 1982; Kleiber 1975), and 17.1 kJ/g (4.09 kcal/g) for glycogen (Kleiber 1975; Whyte et al. 1982). These values represent retained gross chemical energy, not dietary metabolizable energy. The protein value is therefore not the dietary Atwater value of 4 kcal/g. Atwater values estimate the metabolizable energy available from ingested nutrients after digestion, absorption, urinary losses, and other losses. The present calculation asks how much chemical energy is stored in newly deposited tissue. For that purpose, gross-energy values are the appropriate accounting basis.

### 2.3 Biochemical synthesis-cost parameters

Biochemical synthesis cost refers to ATP or ATP-equivalent expenditure used to assemble molecules. For protein, we used 5 ATP equivalents per amino acid incorporated into protein, following the convention used in tissue-deposition modelling (Hall 2010). This value is slightly more inclusive than the minimal translational cost, commonly counted as approximately 4 high-energy phosphate equivalents per incorporated amino acid, because it includes the ATP-equivalent cost assigned to amino acid incorporation in deposition models rather than only ribosomal elongation.

The protein synthesis-cost parameter was derived as follows. One mole of incorporated amino acid residues was assigned 5 mol ATP equivalents. Each mole of ATP was assigned an effective free-energy equivalent of approximately 80 kJ/mol ATP. Thus, one mole of incorporated residues costs approximately 5 x 80 = 400 kJ. Using an average amino acid residue mass of approximately 110 g/mol gives 400 kJ / 110 g = 3.6 kJ/g protein (0.86 kcal/g protein).

For intramuscular triacylglycerol, we used 0.75 kJ/g (0.18 kcal/g), following Hall’s lower-bound estimate for the direct energetic cost of lipid deposition (Hall 2010). This is consistent with treating the small intramuscular triacylglycerol pool as being deposited mainly from preformed fatty acids, rather than assuming substantial de novo lipogenesis from carbohydrate (Hellerstein 1999; Acheson et al. 1988).

For glycogen, we assigned one ATP-equivalent cost per glucose residue. Using 80 kJ/mol and a glucose-residue mass of 162 g/mol gives approximately 0.49 kJ/g (0.12 kcal/g). Even assigning two ATP equivalents would increase the glycogen term by only approximately 10 kJ (2 kcal) per kg wet muscle and would not affect the rounded estimate.

### 2.4 Physiological deposition-cost parameters

#### 2.4.1 Protein deposition efficiency

Effective protein deposition efficiency (kP) is an input parameter. It denotes the fraction of metabolizable energy assigned to protein deposition that is ultimately retained as new skeletal-muscle protein. The cost is then computed from the retained protein energy and kP. This parameter is quantitatively dominant because protein accounts for most of the retained chemical energy in wet skeletal muscle. It is also biologically complex. The metabolizable energy required to retain muscle protein in vivo includes amino-acid activation and peptide-bond formation, but also active amino-acid transport, folding and quality control, breakdown and resynthesis, myofibrillar remodeling, extracellular-matrix remodeling, and heat dissipation. Therefore, kP is not a molecular constant. It is a whole-system coefficient that implicitly absorbs turnover, remodeling, quality control, and other processes required for net muscle-protein retention.

Human studies have defined several adjacent quantities, including post-exercise muscle protein synthesis and breakdown, net amino-acid or protein balance after essential amino-acid ingestion, dose-dependent and distribution-dependent stimulation of myofibrillar protein synthesis by protein feeding, and the effect of protein supplementation on resistance-training-induced gains in fat-free mass and muscle size (Phillips et al. 1997; Tipton et al. 1999; Moore et al. 2009; Areta et al. 2013; Morton et al. 2018). However, these studies do not provide a direct coefficient for the fraction of metabolizable energy retained as new skeletal-muscle protein during adult hypertrophy.

To our knowledge, there are no direct data on protein deposition efficiency in adult humans during skeletal-muscle accretion. The available human evidence comes mainly from infants, which is valuable because it is human, but limited because infants are in a developmental whole-body growth state rather than an adult muscle-remodeling state. If anything, adult skeletal-muscle protein deposition efficiency may be lower than infant whole-body protein deposition efficiency because adult hypertrophy is slower, more localized, and more dependent on remodeling. Across the five primary studies that directly measured or calculated protein deposition efficiency, Roberts and Young (1988), Towers et al. (1997), Pullar and Webster (1977), Coyer et al. (1987), and van Milgen, Noblet and Dubois (2001), central values range from 42.0% to 52.0% (Table 1). We therefore used 46% as a pragmatic central value, approximately corresponding to the median of the available estimates, with 42% to 52% as the primary sensitivity range.

**Table 1.** Protein deposition efficiency estimates used in the calculation. Included studies are primary empirical or model-derived estimates that calculated protein deposition efficiency from energy balance, calorimetry, nitrogen balance, protein storage, or nutrient utilization modelling. Converted protein efficiency is the retained energy in deposited protein divided by the total energy assigned to protein deposition, expressed as a percentage.

| Source | Model | How efficiency was derived | Protein efficiency used |
| --- | --- | --- | --- |
| Roberts & Young 1988 | Human infants | Calculated from a human infant energy deposition model | 42.0% |
| Towers et al. 1997 | Low-birth-human infants | Regression using energy intake/expenditure/nitrogen balance/protein storage | 50.5% (46.0 to 56.0%) |
| Pullar & Webster 1977 | Zucker rats | Regression derived from direct calorimetry and nitrogen balance | 44.4% |
| Coyer et al. 1987 | Young rats | Calculated from heat production and growth cost modelling | 45.6% (43.5 to 47.6%) |
| van Milgen et al. 2001 | Growing pigs | Model estimated from indirect calorimetry and nutrient utilization modelling | 52.0% |

#### 2.4.2 Lipid and glycogen deposition efficiencies

Lipid and glycogen are smaller terms than protein, but they still require explicit assumptions. For lipid, we used a fixed deposition efficiency of 82%, calculated as the mean of three available estimates of fat-deposition efficiency: 73.5% from Pullar and Webster (1977), 85.5% from Roberts and Young (1988), and 88.3% from van Milgen et al. (2001).

For glycogen, we used a fixed deposition efficiency of 95%. This value was derived from simple energetic accounting rather than from direct empirical deposition studies. In glycogen synthesis, glucose is incorporated through UDP-glucose, which is formed from glucose-1-phosphate and UTP, with pyrophosphate hydrolysis helping to drive the reaction forward (Nelson and Cox 2021).

Assigning one ATP-equivalent cost per glucose residue gives an estimated efficiency of 97.2%, while assigning two or 2.5 ATP-equivalents to allow for auxiliary costs gives efficiencies of 94.5% and 93.3%, respectively. The mean of these three values is 95%.

### 2.5 Resting-maintenance parameters

The deposition estimate refers to the energetic cost of tissue construction and retention. It does not include the resting energy required to maintain newly accumulated muscle during the period in which it is being gained. This additional maintenance component was therefore calculated separately. The base calculation assumes that 1 kg of wet skeletal muscle is accrued gradually over approximately 84 days, broadly consistent with human resistance-training studies showing approximately 1 kg-scale gains in fat-free mass over 12 to 13 weeks (Morton et al. 2018). If accretion is approximately linear, the average additional muscle mass present during the accretion period is 0.5 kg. Wang et al. (2010) reported a tissue-specific resting metabolic rate for skeletal muscle of approximately 54.4 kJ/kg/day (13 kcal/kg/day).

### 2.6 Diet-induced thermogenesis parameter

The expanded biological cost must be translated into additional metabolizable energy intake. Additional food intake is accompanied by diet-induced thermogenesis, because part of the ingested metabolizable energy is dissipated as heat during digestion, absorption, transport, metabolism, and storage (Westerterp 2004). We approximated diet-induced thermogenesis as 10% of additional metabolizable energy intake. Therefore, 90% of the additional intake was assumed to remain available to cover the expanded biological cost after deposition and resting maintenance. The required additional metabolizable energy intake was calculated by dividing this expanded biological cost by 0.90

## 3 CALCULATIONS

### 3.1 Component 1 — Stored tissue energy

Stored tissue energy is the chemical energy contained in the newly deposited muscle tissue. For the reference composition:

Protein: 177 g x 23.5 kJ/g = 4159.5 kJ (994 kcal)

Lipid: 30 g x 38.9 kJ/g = 1167 kJ (279 kcal)

Glycogen: 20 g x 17.1 kJ/g = 342 kJ (82 kcal)

(1) Total stored tissue energy = 4159.5 + 1167 + 342 = 5668.5 kJ/kg (1355 kcal/kg).

### 3.2 Component 2 — Biochemical synthesis cost

Biochemical synthesis cost is the lower-bound ATP-equivalent cost of assembling the protein, lipid, and glycogen contained in the new tissue:

Protein synthesis cost: 177 g x 3.6 kJ/g = 637.2 kJ (152 kcal)

Triacylglycerol synthesis cost: 30 g x 0.75 kJ/g = 22.5 kJ (5 kcal)

Glycogen synthesis cost: 20 g x 0.49 kJ/g = 9.8 kJ (2 kcal)

(2) Total biochemical synthesis cost = 637.2 + 22.5 + 9.8 = 669.5 kJ/kg (160 kcal/kg).

Stored tissue energy plus biochemical synthesis cost = 5668.5 + 669.5 = 6338 kJ/kg (1515 kcal/kg).

### 3.3 Component 3 — Physiological deposition cost

Physiological deposition cost is the additional metabolizable energy required because tissue deposition in vivo is not perfectly efficient. It reflects the physiological cost of retaining protein, lipid, and glycogen in new skeletal muscle beyond the stored tissue energy and biochemical synthesis cost already calculated in Sections 3.1 and 3.2.

The total physiological deposition requirement was calculated by dividing the retained energy of each retained substrate by its deposition efficiency:

Physiological deposition requirement = (4159.5 / kP) + (1167 / 0.82) + (342 / 0.95) kJ/kg. Using the central kP value of 0.46:

Physiological deposition requirement = (4159.5 / 0.46) + (1167 / 0.82) + (342 / 0.95) = 10826 kJ/kg (2587 kcal/kg).

Component 3 was then calculated as the difference between this physiological deposition requirement and the biochemical lower bound:

(3) Physiological deposition cost = 10826 − 6338 = 4487.9 kJ/kg (1072 kcal/kg).

Thus, after Component 3, the central running total is 10830 kJ/kg (2587 kcal/kg). Using the primary kP sensitivity range, the physiological deposition requirement is 9780 kJ/kg (2338 kcal/kg) at kP = 52%, 10830 kJ/kg (2587 kcal/kg) at kP = 46%, and 11690 kJ/kg (2793 kcal/kg) at kP = 42%.

### 3.4 Component 4 — Resting maintenance during accretion

Resting maintenance during accretion is the resting energy required to maintain newly accumulated muscle while the 1 kg of tissue is being gained. Under the 84-day linear-accretion assumption, the average additional muscle mass present during the accretion period is 0.5 kg. Therefore:

(4) Resting maintenance = 54.4 kJ/kg/day x 0.5 kg x 84 days = 2285 kJ/kg (546 kcal/kg).

Adding this component gives an expanded biological cost of 10826 + 2285 = 13111 kJ/kg (2587 + 546 = 3133 kcal/kg). Across the primary kP range, the expanded biological cost is 12070 to 13970 kJ/kg (2884 to 3339 kcal/kg).

### 3.5 Component 5 — Diet-induced thermogenesis

Diet-induced thermogenesis is the intake-level cost of processing the additional food required to cover deposition and resting maintenance. Because diet-induced thermogenesis was approximated as 10% of additional metabolizable energy intake, 90% of the additional intake was assumed to remain available to cover the expanded biological cost. Therefore: Additional metabolizable energy intake = expanded biological cost / 0.90.

For the central estimate:

13111 kJ/kg / 0.90 = 14568 kJ/kg (3133 kcal/kg / 0.90 = 3481 kcal/kg).

(5) The diet-induced thermogenesis component is therefore 14568 - 13111 = 1457 kJ/kg (348 kcal/kg).

Across the primary kP range, the final additional metabolizable energy intake is 13410 to 15520 kJ/kg (3204 to 3710 kcal/kg), with a central estimate of 14570 kJ/kg (3481 kcal/kg) (Figure 1).

**Figure 1.**
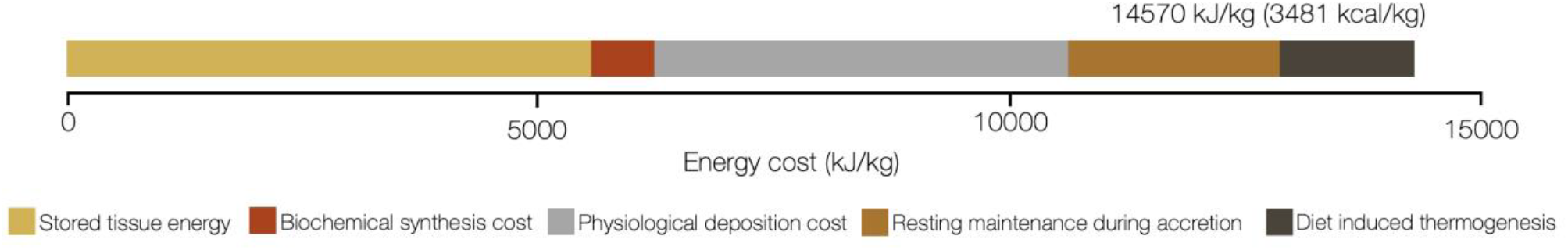
Five-component decomposition of the estimated energetic requirement for building 1 kg of wet human skeletal muscle, expressed as the equivalent additional metabolizable energy intake. Values are central rounded estimates.

### 4. EMPIRICAL TRIANGULATION

The preceding calculation is theoretical, although it is grounded in empirical estimates of stored tissue energy, biochemical synthesis cost, deposition-efficiency parameters, resting maintenance, and diet-induced thermogenesis. Because no study has directly measured the energy required to build 1 kg of adult human skeletal muscle, the estimate requires triangulation against the closest available independent empirical evidence. Studies used to derive the model coefficients were not reused as validation evidence, to avoid non-independent triangulation.

Slater et al. (2019) deserve credit for explicitly recognizing that this quantity was missing and for providing a useful first framework for thinking about the energy cost of skeletal-muscle hypertrophy. The human estimates summarized by Slater et al. (2019), 7440 kJ/kg (1778 kcal/kg) and 6050 kJ/kg (1446 kcal/kg), are useful lower-bound comparators. They are close to the present stored-plus-synthesis estimate of 6340 kJ/kg (1515 kcal/kg). This supports the interpretation that those values approximate stored tissue energy plus limited synthetic overhead rather than the full physiological or intake-level cost of building 1 kg of adult human skeletal muscle.

### 4.1 Independent lean-tissue accumulation estimates in growing rats

The most relevant independent animal comparators are two recent studies that estimated the energetic cost of lean-tissue accumulation in growing rats.

Sekiguchi et al. (2024) estimated the energy expended to synthesize lean tissue in rats at approximately 12.1 kJ/g, equivalent to 2.9 kcal/g. Using their stored lean-tissue energy value of approximately 1.25 kcal/g, equivalent to 5.23 kJ/g, the total energy associated with lean-tissue accumulation becomes:

12.1 kJ/g + 5.23 kJ/g = 17.4 kJ/g, or 17400 kJ/kg lean tissue (4150 kcal/kg lean tissue).

Obikawa et al. (2025) used a related regression approach in growing rats and estimated the energy expended to synthesize lean tissue at 13.9 kJ/g. Adding the same stored lean-tissue energy value gives:

13.9 kJ/g + 5.23 kJ/g = 19.1 kJ/g, or 19100 kJ/kg lean tissue (4570 kcal/kg lean tissue).

Together, these studies suggest an empirical lean-tissue accumulation cost of approximately 17400 to 19100 kJ/kg lean tissue (4150 to 4570 kcal/kg). These values are higher than the present central estimate of 14570 kJ/kg wet skeletal muscle (3481 kcal/kg), but they are of the same order of magnitude. The difference is expected because the rat studies estimate whole-body lean-tissue accumulation, not isolated adult skeletal-muscle hypertrophy. Whole-body lean tissue includes skeletal muscle, organs, skin, connective tissue, bone-associated organic matrix, extracellular components, and other non-fat tissues. Regression-derived lean-tissue coefficients may also absorb broader growth-associated costs rather than only the cost of depositing muscle protein, lipid, glycogen, and water.

### 4.2 Cultured meat as an engineered-system comparator

Tuomisto and Teixeira de Mattos (2011) estimated total primary energy use for cultured meat production at 26000 to 33000 kJ/kg. Using the midpoint of this range, 29500 kJ/kg, and their estimate that the muscle-cell cultivation process accounts for approximately 72% of total energy use, the cultivation component is:

29500 kJ/kg x 0.72 = 21200 kJ/kg (5076 kcal/kg).

This value should not be interpreted as a biological ATP cost or as an in vivo metabolizable-energy cost because it includes ex vivo bioreactor requirements and primary energy inputs. However, it provides a conservative engineered-system upper comparator for the energy required to produce 1 kg of skeletal-muscle tissue under tightly controlled conditions. Its value is higher than the present central intake-level estimate, as expected, because the cultured-meat estimate includes system-level production costs that are absent from in vivo human muscle accretion.

## 5. FROM TISSUE ENERGETICS TO DIETARY SURPLUS

If 1 kg of wet skeletal muscle is accrued over approximately 84 days, corresponding to a 12-week resistance-training period (Morton et al. 2018), the central final estimate of 14570 kJ/kg (3481 kcal/kg) corresponds to approximately 170 kJ/day (41 kcal/day). Across the primary kP range, the daily value is approximately 160 to 190 kJ/day (38 to 44 kcal/day).

This daily value should not be interpreted as the total energetic cost of a hypertrophy intervention. The calculation estimates the additional metabolizable energy intake required to support construction and maintenance of the newly accumulated skeletal-muscle tissue itself. It does not include the energy cost of the resistance-training sessions used to induce hypertrophy, nor broader systemic costs of training adaptation, repair, inflammation, connective-tissue remodeling, or changes in other tissues. The value is therefore a tissue-specific energetic coefficient, not a full intervention-level energy budget.

This distinction matters when the estimate is compared with common sports-nutrition recommendations. Daily energy surpluses of approximately 1260 to 2090 kJ/day (300 to 500 kcal/day) are often suggested during resistance-training phases (Larson-Meyer et al. 2022; Slater et al. 2019). Over 84 days, the midpoint of this range, 1670 kJ/day (400 kcal/day), corresponds to 140600 kJ (33,600 kcal) of additional dietary energy. By contrast, the central calculated additional metabolizable energy intake required to build 1 kg of wet skeletal muscle is 14570 kJ (3481 kcal). The common applied surplus is therefore almost tenfold higher than the calculated tissue-specific requirement.

This discrepancy does not necessarily mean that applied surpluses are irrational. Traditional surplus recommendations may be pragmatic and conservative weight-gain prescriptions designed to reduce the risk of underfeeding in applied settings. They may also cover training expenditure, uncertainty in maintenance-energy estimation, dietary adherence, adaptive changes in energy expenditure, glycogen restoration, recovery demands, and tolerance for some fat gain. However, they should not be interpreted as estimates of the energy required to deposit skeletal muscle itself, because intervention studies show that higher or faster energy surpluses can increase fat mass without proportional gains in skeletal muscle (Garthe et al. 2013; Helms et al. 2023; Sanchez et al. 2024; Slater et al. 2019).

The practical message is therefore not that a resistance-training athlete should simply add 170 kJ/day (41 kcal/day). The estimate is too small relative to the normal uncertainty of applied energy prescription to be used in that way. Its value lies in serving as an austere physiological reference point. It imposes a quantitative boundary on claims about muscle accretion and leaves little room for assuming that large daily surpluses primarily reflect the energy cost of skeletal-muscle deposition itself.

## 6. LIMITATIONS

This analysis has several limitations.

First, the estimate depends on assumed skeletal-muscle composition. Protein, lipid, and glycogen contents vary by muscle group, fiber-type distribution, training status, diet, sex, age, metabolic state, and recent exercise. The use of 177 g protein, 30 g lipid, and 20 g glycogen per kg wet skeletal muscle should therefore be interpreted as a plausible mixed-muscle estimate, not as a universal biological constant (Stroh et al. 2021; Goodpaster et al. 2004; Knuiman et al. 2015; Koopman et al. 2006).

Second, the protein-deposition efficiency assumption dominates the physiological estimate. The central value used here, 46%, with a range of 42 to 52%, is derived from the best available primary evidence, but mainly from infants and growing animals rather than adult humans. These data provide explicit energetic-efficiency coefficients, but they are imperfect analogues of adult skeletal-muscle accretion, which is likely slower, more localized, and more dependent on remodeling than developmental growth. Thus, kP remains the main uncertainty in the model.

Third, the model does not explicitly quantify each biological process that accompanies skeletal-muscle tissue accretion. Satellite-cell activation, myonuclear accretion, connective-tissue remodeling, capillarization, extracellular-matrix turnover, mitochondrial remodeling, inflammation, repair, and expansion of supporting cellular and extracellular structures may all contribute to the energetic cost of building new skeletal muscle. In the present framework, these processes are included only to the extent that they are embedded within kP or the additional resting-maintenance component.

Fourth, the resting-maintenance and diet-induced thermogenesis terms depend on simplifying assumptions. The 84-day accretion period and linear-accretion model provide a practical reference case, not a universal biological trajectory. Similarly, the 10% diet-induced thermogenesis assumption is a reasonable approximation for intake-level accounting, but the true value varies with macronutrient composition, energy balance, and individual physiology.

Fifth, the estimate has not been directly measured in adult humans. Experimental validation would be valuable for testing and refining the present model, but theory and experiment have complementary epistemic roles: quantitative modelling can make explicit biologically meaningful quantities that are difficult to isolate directly from empirical data alone (Phillips 2024). A definitive experimental test would require controlled energy intake, accurate measurement of total energy expenditure, high-resolution assessment of skeletal-muscle mass, preferably by magnetic resonance imaging or computed tomography, repeated body-composition measurements, and correction for changes in fat mass, glycogen, water, extracellular fluid, and non-muscle lean tissues over a period of verified net muscle gain. Until such data exist, the present value should be regarded as a physiologically grounded quantitative estimate with explicit and testable assumptions.

## 7. CONCLUSION

The present analysis estimates the energetic cost of building 1 kg of wet human skeletal muscle. The calculation is organized around five components: stored tissue energy, biochemical synthesis cost, physiological deposition cost, resting maintenance during accretion, and diet-induced thermogenesis. Under the present assumptions, the final additional metabolizable energy intake required to build 1 kg of wet skeletal muscle is approximately 13410 to 15520 kJ/kg (3204 to 3710 kcal/kg), with a central estimate of 14570 kJ/kg (3481 kcal/kg).

This value is greater than stored tissue energy plus direct biochemical synthesis cost, but far smaller than the cumulative energy surplus implied by many applied weight-gain prescriptions. It should not be used as a simple daily dietary prescription. Its role is to provide a tissue-specific physiological reference point against which claims about body mass gain, lean mass gain, dietary surplus, and skeletal-muscle deposition can be judged. By making this previously implicit quantity explicit, the analysis provides an integrated quantitative reference estimate for the energetics of human skeletal-muscle building.

### Declaration of generative AI and AI-assisted technologies in the writing process

During the preparation of this manuscript, the authors used ChatGPT (OpenAI) to assist with language refinement, manuscript organization, and the clear presentation of calculations and scientific arguments. All AI-assisted content, calculations, and references were critically reviewed and independently verified by the authors. The authors take full responsibility for the content of the manuscript.

## Supporting information

Supplementary_Data_1_Energetic_Cost_Model

